# Whole-genome fingerprint of the DNA methylome during chemically induced differentiation of the human AML cell line HL-60/S4

**DOI:** 10.1101/608695

**Authors:** Enoch Boasiako Antwi, Ada Olins, Vladimir B Teif, Matthias Bieg, Tobias Bauer, Zuguang Gu, Benedikt Brors, Roland Eils, Donald Olins, Naveed Ishaque

**Affiliations:** Division of Theoretical Bioinformatics, German Cancer Research Center (DKFZ), Heidelberg, Germany; Molecular and Cellular Engineering, Centre for Biological Signalling Studies, Freiburg University, Germany; Department of Pharmaceutical Sciences, College of Pharmacy, University of New England, Portland, ME, USA; School of Biological Sciences, University of Essex, Colchester, UK; Germany Heidelberg Center for Personalized Oncology (DKFZ-HIPO), German Cancer Research Center (DKFZ), Heidelberg, Germany; Division of Applied Bioinformatics, German Cancer Research Center (DKFZ), Heidelberg, Germany; Center for Digital Health, Berlin Institute of Health and Charité - Universitätsmedizin Berlin, Kapelle-Ufer 2, 10117, Berlin, Germany; Translational Lung Research Center Heidelberg (TLRC), German Center for Lung Research (DZL), University of Heidelberg, Heidelberg, Germany

**Author notes:** Corresponding Author: Naveed Ishaque.

**Keywords:** DNA Methylation, Promyelocyte, Granulocyte, Macrophage, Differentiation, Epigenetics, Enhancer, Promoter, Multi-omics correlation

## Abstract

**Background:** Myeloid differentiation gives rise to a plethora of immune cells in the human body. This differentiation leaves strong signatures in the epigenome through each differentiated state of genetically identical cells. The leukemic HL-60/S4 promyelocytic cell can be easily differentiated from its undifferentiated promyelocyte state into neutrophil-and macrophage-like cell states, making it an excellent system for studying myeloid differentiation. In this study, we present the underlying genome and epigenome architecture of HL-60/S4 through its undifferentiated and differentiated cell states.

**Results:** We performed whole genome bisulphite sequencing of HL-60/S4 cells and their differentiated counterparts. With the support of karyotyping, we show that HL-60/S4 maintains a stable genome throughout differentiation. Analysis of differential CpG methylation reveals that most methylation changes occur in the macrophage-like state. Differential methylation of promoters was associated with immune related terms. Key immune genes, CEBPA, GFI1, MAFB and GATA1 showed differential expression and methylation. However, we observed strongest enrichment of methylation changes in enhancers and CTCF binding sites, implying that methylation plays a major role in large scale transcriptional reprogramming and chromatin reorganisation during differentiation. Correlation of differential expression and distal methylation with support from chromatin capture experiments allowed us to identify putative proximal and long-range enhancers for a number of immune cell differentiation genes, including CEBPA and CCNF. Integrating expression data, we present a model of HL-60/S4 differentiation in relation to the wider scope of myeloid differentiation.

**Conclusions:** For the first time, we elucidate the genome and CpG methylation landscape of HL-60/S4 during differentiation. We identify all differentially methylated regions and positions. We link these to immune function and to important factors in myeloid differentiation. We demonstrate that methylation plays a more significant role in modulating transcription via enhancer reprogramming, rather than by promoter regulation. We identify novel regulatory regions of key components in myeloid differentiation that are regulated by differential methylation. This study contributes another layer of “omics” characterisation of the HL-60/S4 cell line, making it an excellent model system for studying rapid *in vitro* cell differentiation.

**Summary statement:** Epigenomics plays a major role in cell identity and differentiation. We present the DNA methylation landscape of leukemic cells during in-vitro differentiation, to add another ‘omics layer to better understand the mechanisms behind differentiation.

## Introduction

Gene expression profiles differ among different cell types and change as stem cells differentiate (Cheng *et al.*, 1996; Le Naour *et al.*, 2001; Natarajan *et al.*, 2012). Genome wide CpG methylation, an epigenetic regulation and modification process, has been shown to exhibit similar dynamic behaviour during differentiation (Brunner *et al.*, 2009; Bock *et al.*, 2012). Usually, these two changes (i.e., gene expression and CpG methylation) have been shown to correlate negatively with each other, depending upon the location of the methylated CpG relative to the gene body (Payer *et al.*, 2008; Chuang, Chen and Chen, 2012; Jones, 2012; Yang *et al.*, 2014). Overall, changes in methylation patterns between cell types and tissues throughout life, work to either activate or shut down specific cellular processes (Smith & Meissner, 2013), making cells exhibit different phenotypic characteristics. Acting as a shutdown mechanism, DNA methylation reinforces gene silencing, when expression is not required in a particular cell type (Lock, et al., 1987).

Normal myeloid cell differentiation occurs within the bone marrow, where stroma cells secrete cytokines to help activate myeloid-specific gene transcription (De Kleer, et al., 2014). Further differentiation can occur in the peripheral tissues or blood, dependent upon exposure of the myeloid precursors to cytokines and other factors, such as antigens (Geissmann *et al.*, 2010; Álvarez-Errico *et al.*, 2015). The first direct committed step toward myeloid cell development is the differentiation of multipotent progenitors (MPP) cells into common myeloid progenitor cells (CMP) (Kondo, et al., 1997), (Alvarez-Errico, et al., 2015). CMP cells can then differentiate further into the granulocyte-macrophage lineage progenitor (GMP) and megakaryocyte-erythroid progenitor (MEP) (Iwasaki & Akashi, 2007). While CMP cells can differentiate into all myeloid cell types, GMP cells give rise mainly to monocytes/macrophages and neutrophils, together with a minor population of eosinophils, basophils and mast cells (Laiosa, et al., 2006), (Iwasaki & Akashi, 2007), (Alvarez-Errico, et al., 2015).

The human myeloid leukemic cell line HL-60/S4 is an excellent system to study epigenetic changes during chemically induced in vitro cell differentiation. HL-60/S4 cells are supposedly blocked at the GMP cell state and unable to differentiate any further. The HL-60/S4 cell line is a subline of HL-60 and demonstrates “faster” cell differentiation than the parent HL-60 cells. Undifferentiated HL-60/S4 cells exhibit a myeloblastic or promyelocytic morphology with a rounded nucleus containing 2 to 4 nucleoli, basophilic cytoplasm and azurophilic granules (Birnie, 1988). Retinoic acid (RA) can induce HL-60/S4 differentiation to a granulocyte-like state. 12-O-tetradecanoylphorbol-13-acetate (TPA) can induce differentiation to monocyte/macrophage-like states (Fontana, Colbert and Deisseroth, 1981; Birnie, 1988).

The extent to which DNA methylation regulates these chemically induced differentiation processes is not known. Likewise, the global genome wide methylation changes associated with these differentiation processes have not been described. This study details the methylation changes (and lack of changes), when HL-60/S4 is differentiated to granulocytes, employing RA, and to macrophage, employing TPA. The information contained within this study is intended as a sequel to previous studies that describe the transcriptomes (Mark Welch *et al.*, 2017), nucleosome positioning (Teif *et al.*, 2017) and epichromatin properties (Olins *et al.*, 2014) of HL-60/S4 cells differentiated under identical conditions. The goal is to integrate these different lines of information into a comprehensive description and mechanistic analysis of the cell differentiation pathways in the human myeloid leukemic HL-60/S4 cell lineage.

## Results

### Little or no DNA methylation changes are observed upon HL-60/S4 cell differentiation at the megabase scale

We performed whole genome bisulphite sequencing (WGBS) of HL-60/S4 in 3 different cell differentiation states: the undifferentiated state (UN), the retinoic acid treated granulocyte state (RA), and the tetradecanoyl phorbol acetate (TPA) treated macrophage state. Comparison of the whole genome coverage profiles for each of the three differentiation states of HL-60/S4 revealed that the cell line is hypo-diploid (Mark Welch, Jauch, Langowski, Olins, & Olins, 2017) and is chromosomally stable throughout differentiation (Supplementary Figure S1 A-C). A comparison of HL-60/S4 cells (from 2008 and 2012) by fluorescent in situ hybridization (FISH) karyotyping showed that this cell line is also stable over long time periods (Supplementary Figure S1 D&E). From all the CpGs identified by WGBS on all three cell states, a total of 21,974,649 (82.38%) CpGs had >= 10x coverage (Table 1 and Table S1), which spanned the full range of methylation rates, from 0 (completed unmethylated) to 1 (fully methylated). Most of these CpGs are highly and fully methylated (> 0.75 methylation rate), with only small sets of lowly and unmethylated CpGs (< 0.25 methylation rate) and partially methylated CpGs (methylation rate from 0.25 to 0.75) (Figure 1 A and B). Principal component analysis of all CpGs with coverage greater than 10 revealed that the RA treated samples differed only slightly from the untreated sample, while the TPA samples had a much higher methylation variance, compared to the other two samples (Figure 1C). However, little or no methylation differences were observed among the 3 samples, when methylation rates were averaged over 10 megabase (Mb) windows (Figure 1D).

**Table 1:**
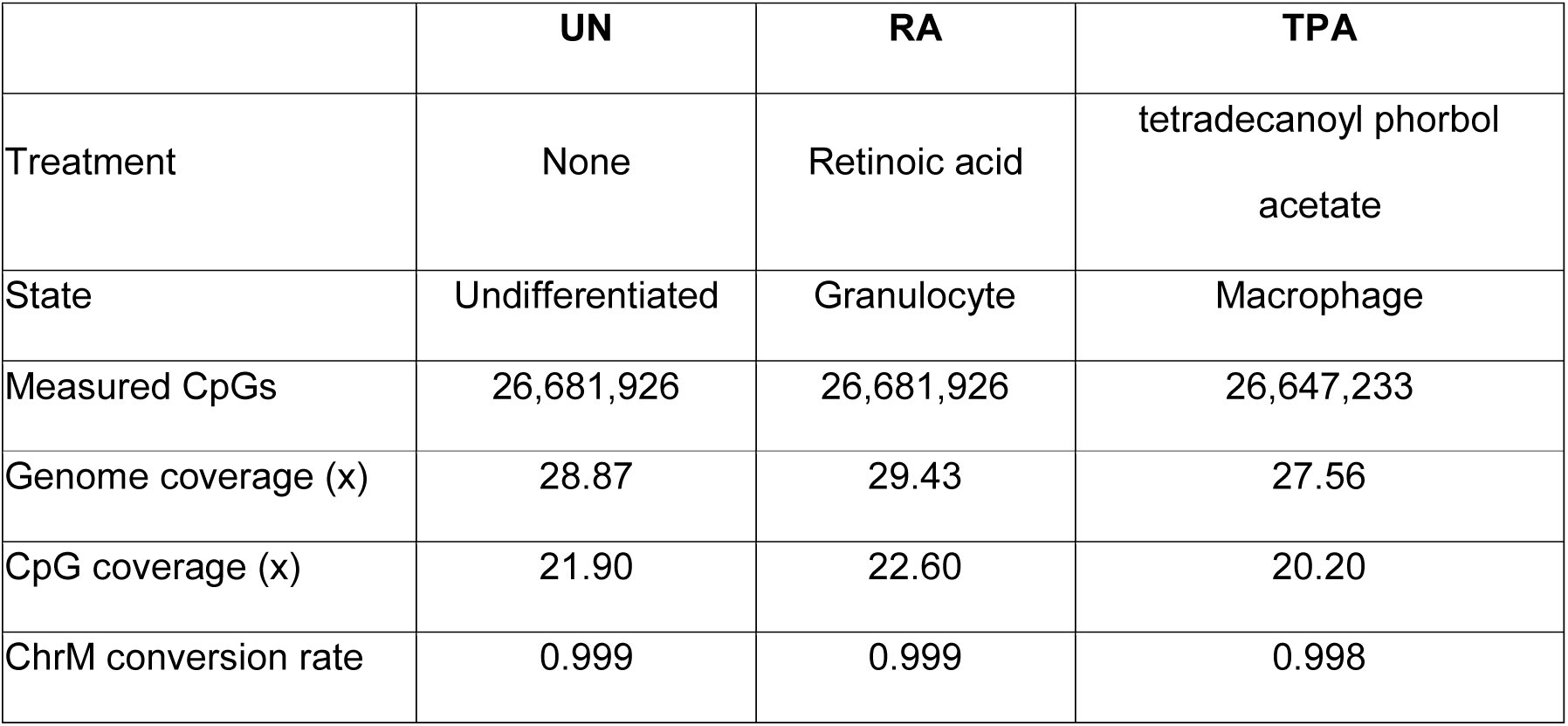
CpG coverage statistics. A summary of the whole genome bisulphite sequencing (WGBS) data for the undifferentiated HL-60/S4 (UN), and retinoic acid (RA) and tetradecanoyl phorbol acetate (TPA) treated cells.

|  | **UN** | **RA** | **TPA** |
| --- | --- | --- | --- |
| Treatment | None | Retinoic acid | tetradecanoyl phorbol<br>acetate |
| State | Undifferentiated | Granulocyte | Macrophage |
| Measured CpGs | 26,681,926 | 26,681,926 | 26,647,233 |
| Genome coverage (x) | 28.87 | 29.43 | 27.56 |
| CpG coverage (x) | 21.90 | 22.60 | 20.20 |
| ChrM conversion rate | 0.999 | 0.999 | 0.998 |

**Figure 1:**
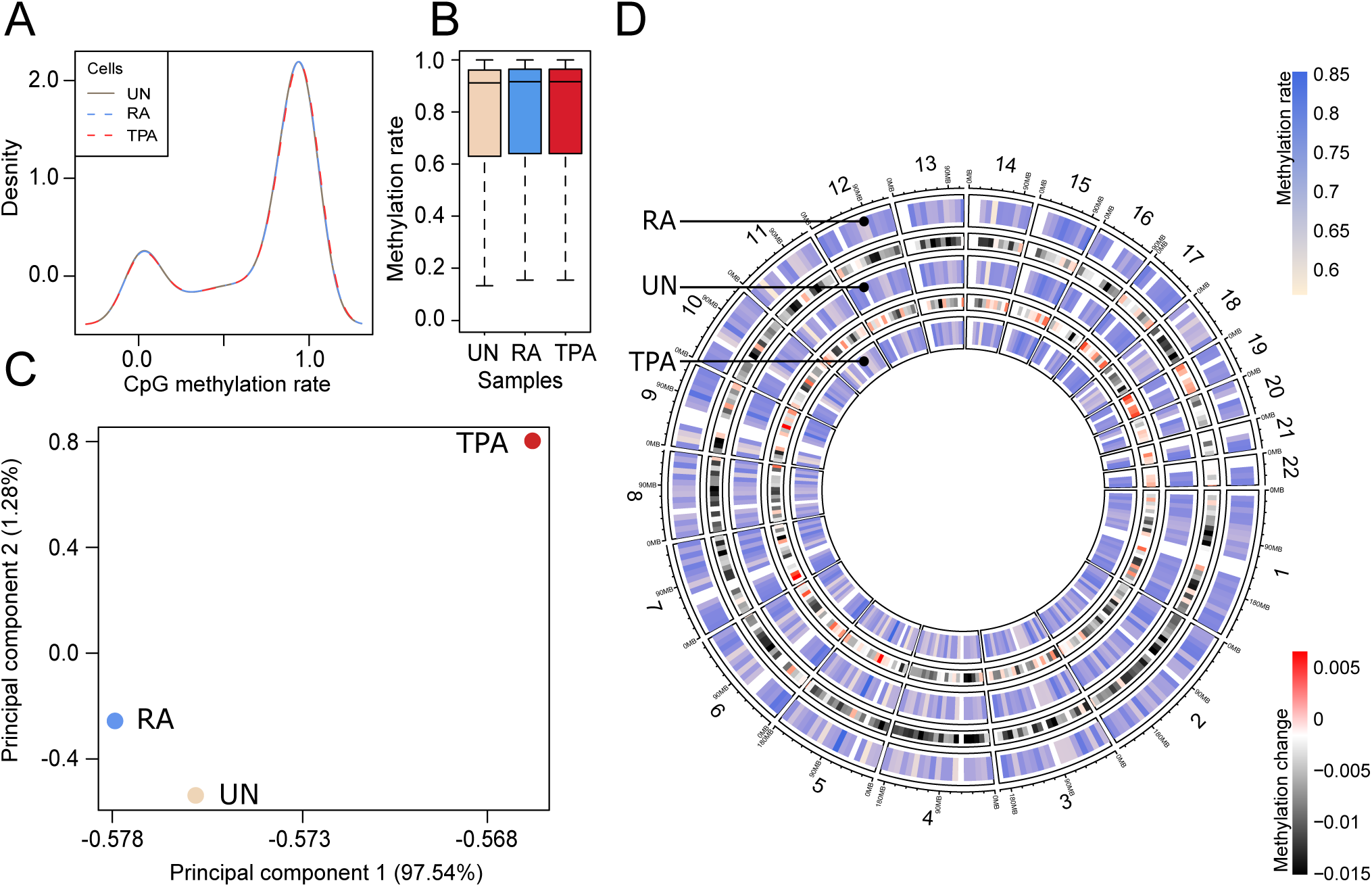
Analysis of DNA methylome upon chemical induction of differentiation of HL-60/S4 cells. A. Whole genome CpG methylation rate density plot. B. Box plots summarising the distribution of CpG methylation rates per sample replicates for the ∼22 million CpGs with coverage greater than or equal to 10x in all samples. The upper and lower limits of the boxes represent the first and third quartiles respectively, and the black horizontal line is the median. The whiskers indicate the variability outside the upper and lower quartiles. C. Principal component analysis of the WGBS data for the three treated samples. D. Circular representation of DNA methylation rates for the different treatments. CpG methylation rates were averaged over 10-Mb windows and are presented as heat map tracks. The heat maps show the DNA methylation change with respect to the sample in the next inner track.

### The single CpG methylation landscape of TPA cells differ most, when compared to UN and RA Cells

Due to the small changes observed on the megabase scale, we focused on significantly differentially methylated single CpGs positions (DMPs) for further analysis. A total of 41,306 unique CpGs were identified to be significantly differentially methylated (Fisher analysis, see Materials and Methods). These DMPs comprise of 12,713, 17,392 and 17,100 CpGs from the comparisons of RA to UN cells, TPA to UN cells and RA to TPA cells, respectively (Figure 2A). A higher proportion of the DMPs identified in the comparison of TPA to UN cells were hyper-methylated; but a similar number of hyper-and hypo-methylated DMPs were observed in the RA to TPA cells comparison. Most of the hyper-methylated DMPs had a methylation rate shift from around 0 to 0.2; hypo-methylated DMPs showed a reverse shift of methylation rate (0.2 to 0) (Figure 2C and D).

**Figure 2:**
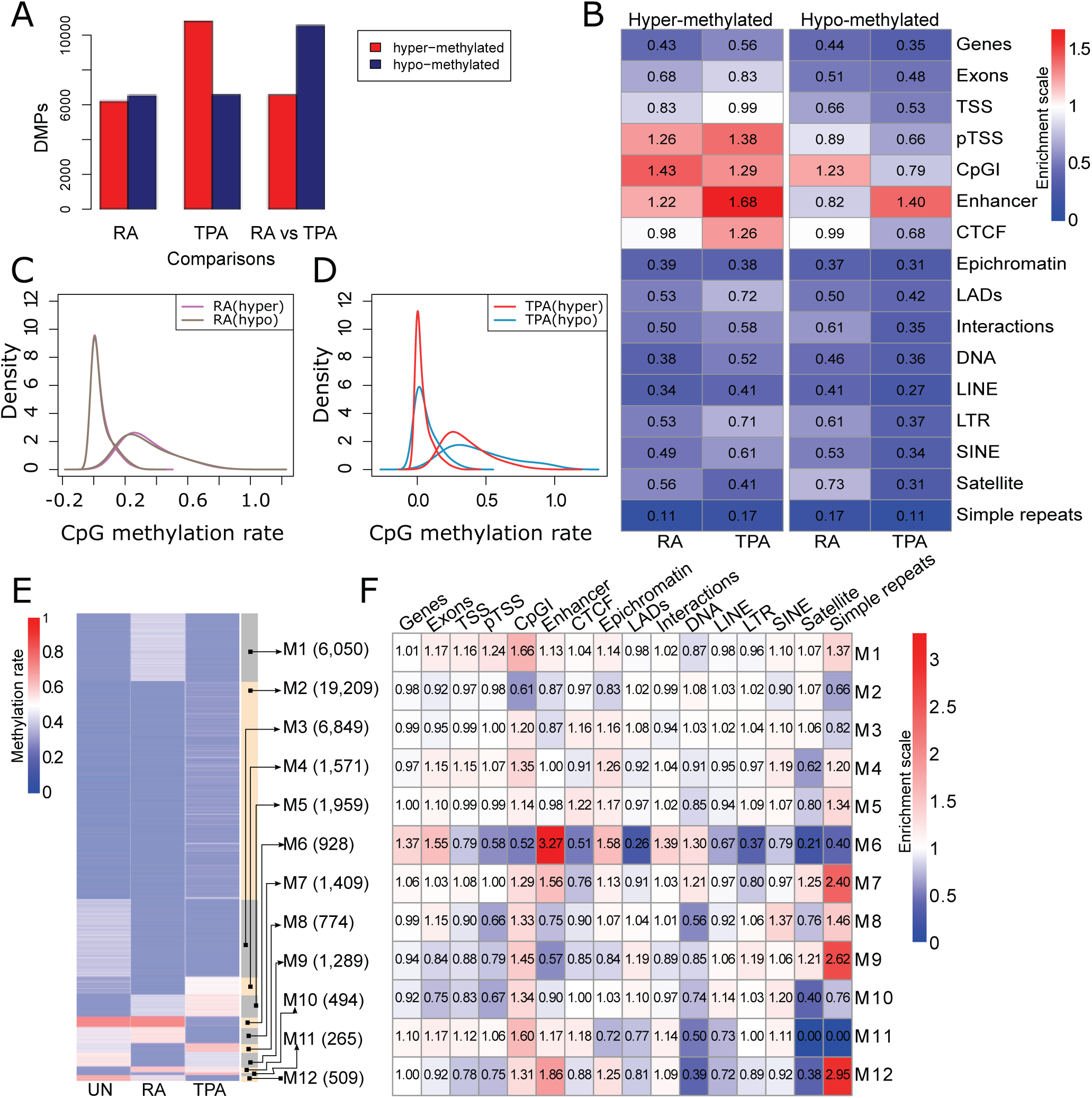
Differentially methylated CpGs (DMPs) analysis. A. Number of DMPs identified with Fisher exact test for each comparison. RA and TPA are the DMPs identified when RA or TPA was compared to the control (UN), while RA vs TPA is the comparison in which RA was compared to TPA. B. Enrichment of genomic features in the hyper-methylated (left) and hypo-methylated (right) DMPs in RA and TPA, compared UN cells. C. The density plot of the methylation rates of DMPs. Hyper and hypo-methylated DMPs are denoted by (hyper) and (hypo) respectively. On the left and right panels show the distribution of DMPs identified in the RA and TPA compared to UN cells respectively. D. Modules identified from the unsupervised clustering of the DMPs. E. Genomic feature enrichment in the 12 modules identified. F. Enrichment of genomic features in the 12 identified modules.

### Enhancers are most enriched within DMPs

The most enriched genomic features in the hyper-methylated DMPs were enhancers, transcription start sites (TSSs) of protein coding genes and CpG islands (CpGIs) for both RA and TPA cells, compared to UN cells (Figure 2B). CTCF was enriched in TPA hyper-methylated DMPs, but not in RA. On the other hand, CpGIs were also the most enriched feature in the hypo-methylated DMPs, when RA was compared to UN cells. Enhancers alone showed a high enrichment in both hyper-methylated and hypo-methylated DMPs, identified when TPA is compared to UN cells (Figure 2B). In contrast to enhancers, simple repeats, epichromatin, and LINE (long interspersed nuclear element) repeats were depleted within hyper-and hypo-methylated regions in both RA and TPA.

We identified clusters of DMP methylation pattern changes between the 3 cell states of HL-60/S4. We called these cluster “modules”. Module analysis reveals that enhancers are significantly enriched in DMPs that are hypo-methylated in the TPA state, relative to UN and RA (modules M6 and M12). The observed hypo-methylation for TPA treated cells corresponded with lower nucleosome occupancy around the DMPs of M6 and M12 (Supplementary Figure S2). M7 DMPs were similarly hypo-methylated in the TPA, compared to RA and UN cells, but with lower methylation differences (Figure 2E and F). Enrichment of exons, epichromatin and chromatin-interacting domains (Li Teng *et al.*, 2015) were also observed in module M6.

### CpGIs have a very dynamic differential methylation

CpGIs are differentially methylated, but mainly in relation to RA treated cells. CpGIs were most enriched in module M1, which has DMPs that are hemi-methylated (approximated 0.5 methylation rate) in RA; but these DMPs showed lower methylation in TPA and UN cells. Similar results were seen in module M9, where DMPs were hypo-methylated in RA, compared to TPA and UN cells. Likewise, CpGI enrichment was observed for module M11, where DMPs are hyper-methylated in TPA, compared to RA and UN cells.

### Methylation of transcription start site DMPs correlate weakly with gene expression

A total of 110 and 132 genes were found to have their TSS overlapping with DMPs from RA and TPA cells compared to UN cells, respectively. These overlapping DMPs had a methylation rate difference of at least 0.2. RA genes showed a weak and insignificant correlation between the average methylation difference of the DMPs overlapping with the TSS and the -log_2_ (RNA expression fold change) of genes (Figure 3A). This is confirmed by the comparable number of genes that have positive and negative correlation between TSS DMP methylation and gene expression (Figure 3B).

**Figure 3:**
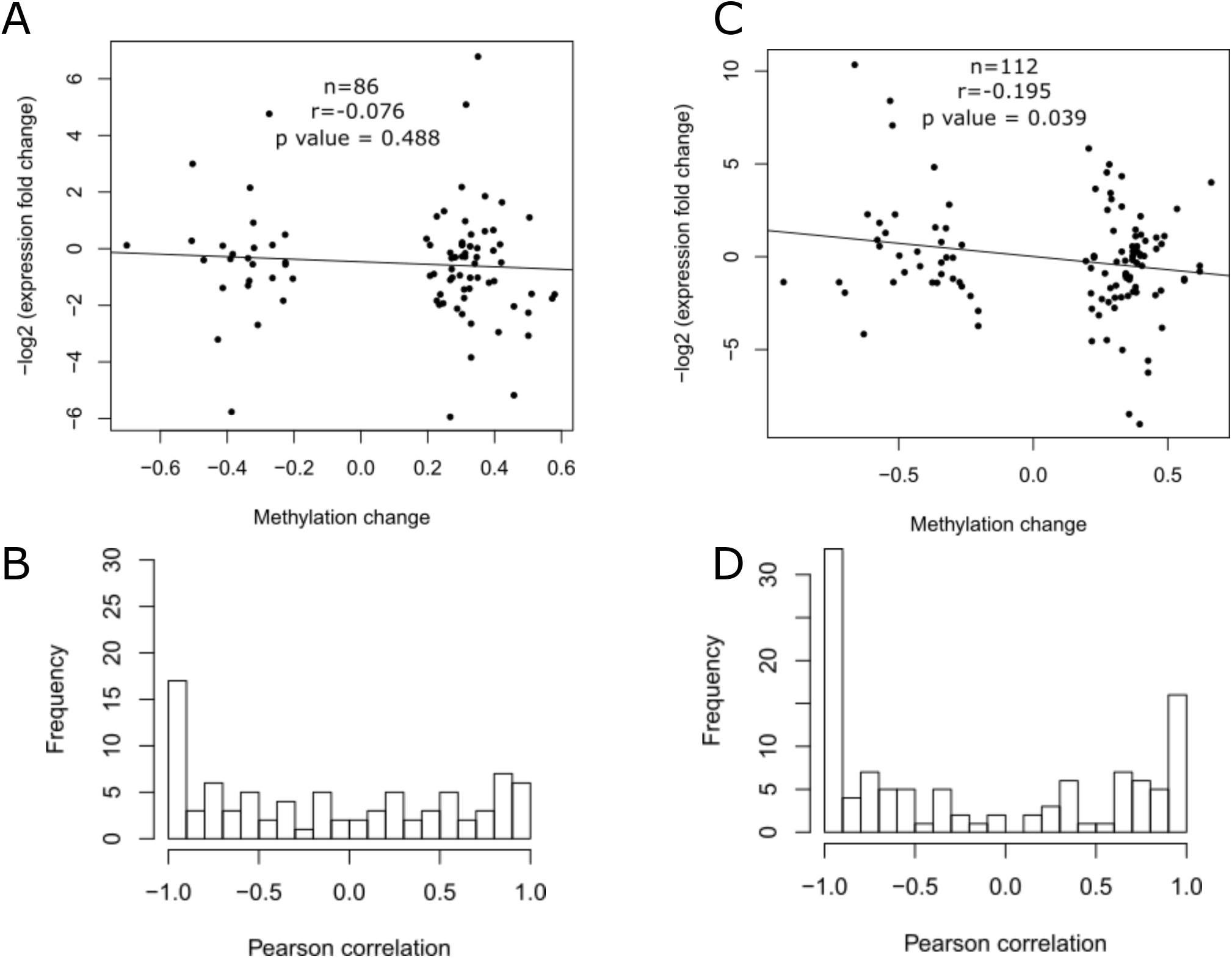
Correlation between TSS methylation and gene expression A. The scatter plot of TSS DMPs methylation change and fold change of RNA expression values (-log2 transformed) for genes which had their TSS overlapping with DMPs with 0.2 methylation difference between RA and UN cells. B. The distribution of Pearson correlation coefficient values between the average methylation of DMPs overlapping with TSS and expression intensity for the genes with differentially methylated TSS in RA relative to UN cells. C. The scatter plot of TSS DMPs methylation change and fold change of expression values (-log2 transformed) for genes which had their TSS overlapping with DMPs with 0.2 methylation difference between TPA and UN cells. D. The distribution of Pearson correlation coefficient values between the average methylation of DMPs overlapping with TSS and expression intensity for the genes with differentially methylated TSS in TPA relative to UN cells.

However, the scatter plot of TSS overlapping DMP methylation change and -log_2_ (RNA expression fold change) does show a weak, but significant, negative correlation (Figure 3C) and a higher number of negatively correlating genes, compared to positively correlating ones (Figure 3D).

### Methylation of long distance regulatory regions shows negative correlation with target gene expression

Methylation and expression of CEBPE (a major transcription factor involved in myeloid cell differentiation) shows a negative correlation at the 3’ end of the gene; a region identified to be an enhancer in the ROADMAP epigenome project (Figure 4A) (Roadmap Epigenomics Consortium *et al.*, 2015). The downstream region of the CEBPE gene, containing the DMPs whose methylation has strong negative correlation with expression, has been shown through IMPET (integrated methods for predicting enhancer targets) and CHIA-PET (Chromatin Interaction Analysis by Paired-End Tag Sequencing) to interact with the upstream regions that spans part of the gene body and the TSS region (L. Teng *et al.*, 2015). No DMPs were observed overlapping the TSS of the CEBPE gene; hence, no correlation between TSS methylation and expression is available. The RNA expression of CEBPE (as well as CCNF and PGP) in UN, RA, and TPA is shown in Figure 4B.

**Figure 4:**
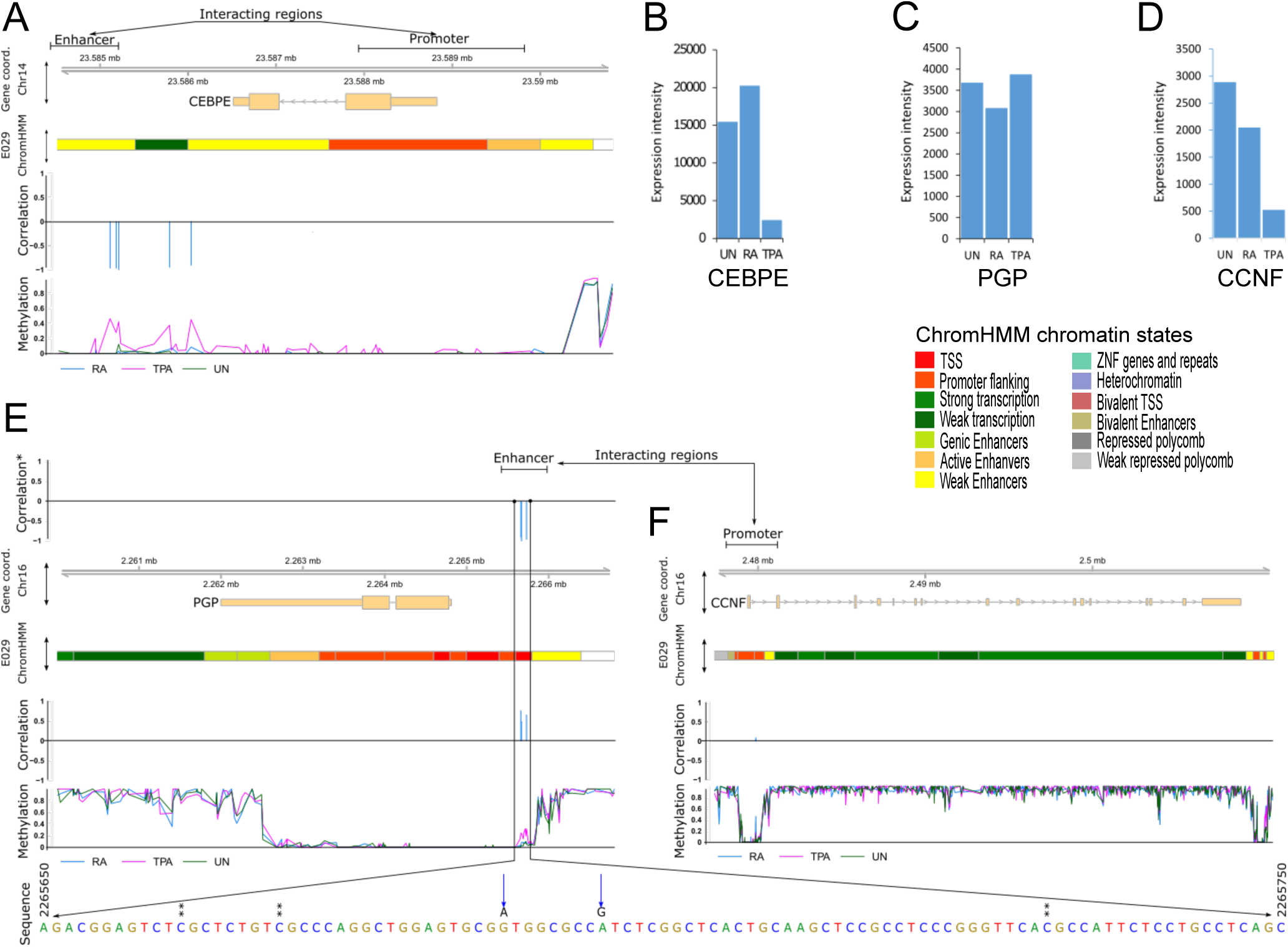
Gene expression integration with CpG differential methylation shows that CEBPE expression is regulated by methylation of downstream region. A. The promoter region of CEBPE gene shows a strong inverse correlation between expression and methylation of DMPs in its downstream region. Top panel: the interacting regions are regions identified with IM-PET in K562 cells and confirmed by CHIA-PET. Second panel shows genomic coordinated of CEBPE gene on chromosome 14 whereas E029 chromHMM panel shows the genomic features along the CEBPE gene in E029 (primary monocyte cells from peripheral blood). Correlation panel shows the correction of between DMPs along the genomic coordinates and the gene expression of CEBPE for all three states. Methylation panel shows the methylation rate of CpGs along the gene coordinates +/-2kb. B-D. The expression of CEBPE (B), PGP (C) and (CCNF) for the three different states. E, F. The cyclin-F-box protein coding gene CCNF interacts with a distant upstream region which regulated its expression through methylation. Panel description is similar to figure 4A except the “correlation*” panel in 4E which depicts the correlation between the methylation of CpGs along the PGP gene coordinates and the expression of CCNF. The sequence panel shows the nucleotide sequence of in the differentially methylated region upstream PGP (enhancer region): DMPs are marked in the sequence panel by ** while the blue arrows points to TPA-specific SNP sites within the differentially methylated region. 4F shows similar information depicted in 4A for CCNF gene. The interaction between the CCNF promoter and the region upstream PGP was also identified by ChIA-PET in the K562 cell line.

Furthermore, the RNA expression of the gene encoding for cyclin F, CCNF, (Figure 4D) correlates weakly with the methylation of DMPs (Figure 4F) that overlap with its gene and TSS. However, CCNF RNA expression has a strong negative correlation with DMPs overlapping the upstream region of the PGP gene, which encodes phophoglycolate phosphatase (Figure 4E correlation*). PGP RNA expression (Figure 4C) does not show a similar correlation. This region has also been identified by ROADMAP as an enhancer.

### Functional annotation of DMPs are mostly immune response related

Using DMPs with a methylation fold change greater than or equal to 2, we observed that immune response related cellular functions were the most enriched biological function for all the genes whose TSS overlapped with DMPs, when RA cells were compared to UN cells (Table 2). Similarly, genes with their TSS overlapping DMPs in TPA compared to UN cells, were also mostly related to (or involved with) phosphoproteins, signalling and defence responses, including chemotaxis (Table 3). Similar observations were made when DMPs were merged into DMRs and their functional associations tested in TPA, compared to UN cells (Table 5). For RA compared to UN cells, the functional annotation was general cell function related (Table 4).

**Table 2:** Immune response related functions are predominant in cellular functions of genes with the most differentially methylated TSS in RA, compared to UN cells. The functional annotation of genes with their TSS overlapping with DMPs, with a methylation rate difference >=0.2 in RA, compared to UN cells, for which gene expression data was available. The p-value is the calculated hypergeometric binomial calculated in DAVID.

| Term | % | Enrichment | PValue |
| --- | --- | --- | --- |
| Glycoprotein binding | 3.85 | 19.86 | 0.01 |
| Translation | 7.69 | 4.46 | 0.01 |
| Defence response | 10.26 | 3.2 | 0.01 |
| Immunoglobulin-like V-type domain | 5.13 | 8.01 | 0.01 |
| Signal peptide | 26.92 | 1.6 | 0.03 |
| Steroid binding | 3.85 | 11.48 | 0.03 |
| Protein biosynthesis | 5.13 | 5.32 | 0.04 |
| Lipoprotein | 8.97 | 2.72 | 0.04 |
| Positive regulation of cell migration | 3.85 | 8.29 | 0.05 |
| Peroxisome | 3.85 | 8.33 | 0.05 |
| Enzyme binding | 7.69 | 2.81 | 0.06 |
| Positive regulation of locomotion | 3.85 | 7.53 | 0.06 |
| Endocytosis signal motif | 2.56 | 27.58 | 0.07 |
| Defence response to Gram-positive bacterium | 2.56 | 24.6 | 0.08 |
| Ankyrin | 5.13 | 3.94 | 0.08 |
| Phospholipid catabolic process | 2.56 | 22.36 | 0.08 |
| Cell membrane | 17.95 | 1.59 | 0.09 |
| SH2 domain binding | 2.56 | 20.41 | 0.09 |
| Locomotory behavior | 5.13 | 3.59 | 0.1 |
| Structure of Caps and SMACs | 2.56 | 14.92 | 0.1 |

**Table 3:** Immune response related functions are predominant in cellular functions of genes with the most differentially methylated TSS in TPA compared to UN cells. The functional annotation of genes with their TSS overlapping with DMPs with a methylation rate difference >=0.2 in TPA, compared to UN for which gene expression data was available. The p-value is the calculated hypergeometric binomial calculated in DAVID.

| **Term** | **%** | **Enrichment** | **PValue** |
| --- | --- | --- | --- |
| Defense response | 11.43 | 3.47 | 0 |
| Phosphatase activity | 5.71 | 4.29 | 0.01 |
| Positive regulation of locomotion | 3.81 | 7.27 | 0.02 |
| Chemotaxis | 5.71 | 3.9 | 0.02 |
| Peroxisome | 3.81 | 7.09 | 0.02 |
| Leukocyte transendothelial migration | 3.81 | 5.75 | 0.03 |
| Opsonization | 1.9 | 59.33 | 0.03 |
| Immunoglobulin-like fold | 7.62 | 2.59 | 0.03 |
| Translation | 5.71 | 3.23 | 0.04 |
| Immune response | 8.57 | 2.32 | 0.04 |
| RNA binding | 8.57 | 2.23 | 0.04 |
| Steroid binding | 2.86 | 8.34 | 0.05 |
| Phosphoprotein | 44.76 | 1.23 | 0.06 |
| Intracellular protein transport | 5.71 | 2.86 | 0.06 |
| p53 signalling pathway | 2.86 | 7.48 | 0.06 |
| MAPK signalling pathway | 4.76 | 3.17 | 0.06 |
| Positive regulation of phagocytosis | 1.9 | 29.67 | 0.06 |
| Regulation of leukocyte activation | 3.81 | 4.29 | 0.06 |
| Zinc finger region:C3H1 | 1.9 | 29.11 | 0.07 |
| Protein biosynthesis | 3.81 | 4.05 | 0.07 |
| Signal peptide | 22.86 | 1.4 | 0.08 |
| Regulation of apoptosis | 8.57 | 1.99 | 0.08 |
| Macrophage activation | 1.9 | 23.73 | 0.08 |
| Small GTPase mediated signal transduction | 4.76 | 2.92 | 0.09 |
| Calcium-binding | 1.9 | 21.03 | 0.09 |
| Antimicrobial | 3.81 | 3.69 | 0.09 |
| Cell cycle | 5.71 | 2.48 | 0.09 |
| Cytoskeleton organization | 5.71 | 2.45 | 0.09 |
| Structure of Caps and SMACs | 1.9 | 14.92 | 0.1 |
| Palmitate moiety binding | 3.81 | 3.58 | 0.1 |

**Table 4:**
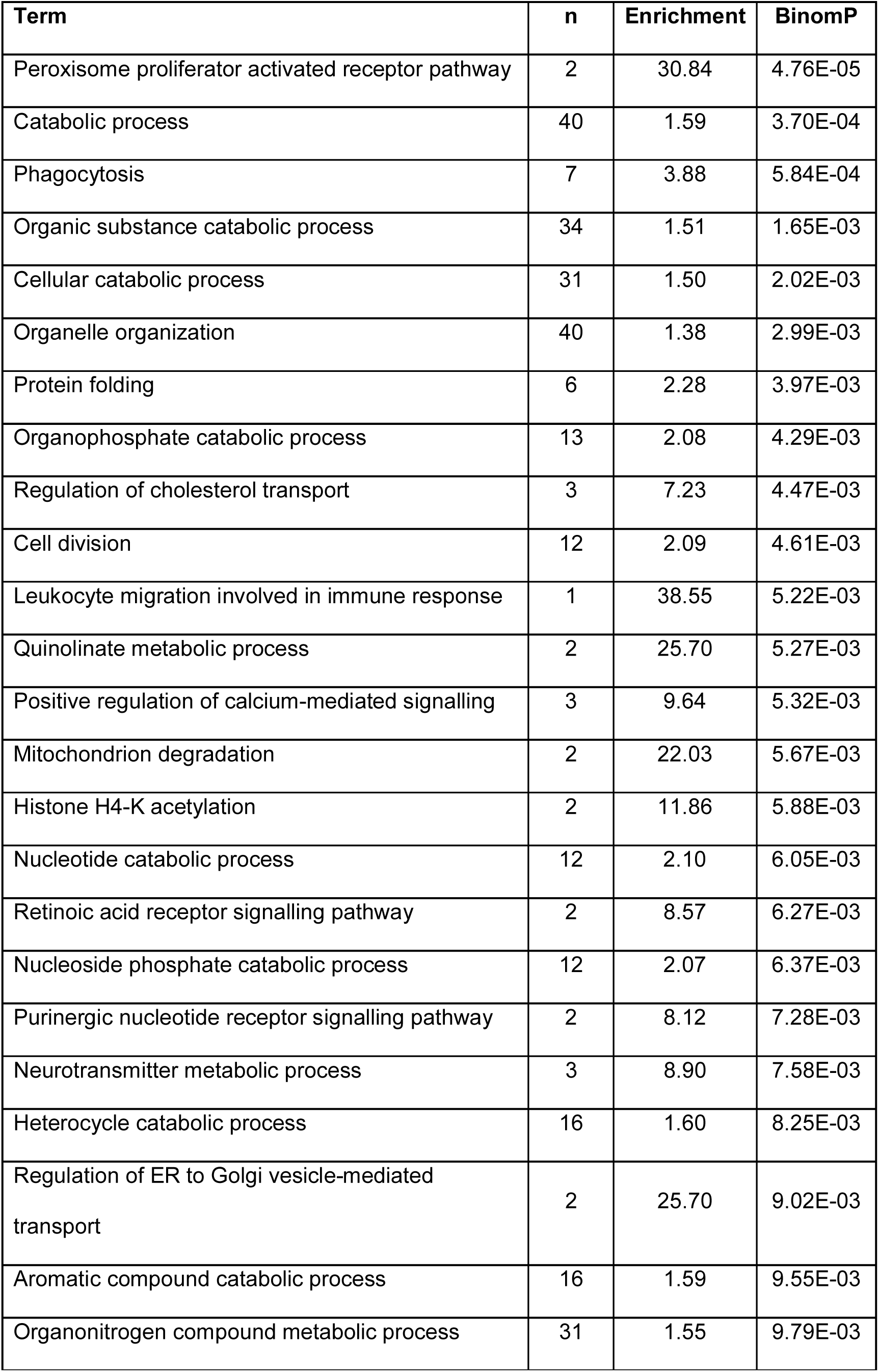
Cellular functions of DMRs which were generated by merging DMPs. The biological process enrichment was performed with DMRs generated from TPA-UN comparison DMPs. The p-value is the calculated binomial calculated by GREAT (McLean *et al.*, 2010).

**Table 5:** Cellular functions of DMRs which were generated by merging DMPs. The biological process enrichment was performed with DMRs generated from TPA-UN comparison DMPs. The p-value is the calculated binomial calculated by GREAT (McLean *et al.*, 2010).

| **Term** | **n** | **Enrichment** | **P value** |
| --- | --- | --- | --- |
| Regulation of defense response | 53 | 1.76 | 1.1E-10 |
| Endocytosis | 50 | 2.14 | 3.8E-10 |
| Negative regulation of interleukin-8 production | 5 | 12.93 | 6.5E-08 |
| Regulation of inflammatory response | 27 | 1.98 | 1.3E-07 |
| Negative regulation of protein modification process | 44 | 1.88 | 1.1E-06 |
| Negative regulation of transferase activity | 27 | 2.15 | 1.1E-06 |
| Immune response-activating signal transduction | 36 | 1.95 | 1.2E-06 |
| Activation of immune response | 38 | 1.75 | 2.4E-06 |
| Positive regulation of defense response | 31 | 2.05 | 3.1E-06 |
| Negative regulation of protein kinase activity | 25 | 2.23 | 3.6E-06 |
| Negative regulation of kinase activity | 26 | 2.19 | 5.7E-06 |
| Negative regulation of phosphorylation | 35 | 2.23 | 8.1E-06 |
| Negative regulation of protein phosphorylation | 34 | 2.30 | 9.0E-06 |
| Platelet activation | 29 | 2.10 | 1.6E-05 |
| Regulation of peptidase activity | 38 | 1.71 | 1.9E-05 |
| Toll-like receptor 5 signalling pathway | 12 | 2.86 | 2.5E-05 |
| Toll-like receptor 10 signalling pathway | 12 | 2.86 | 2.5E-05 |
| Response to lipoprotein particle stimulus | 5 | 7.76 | 6.5E-05 |
| Positive regulation of histone H4 acetylation | 3 | 15.51 | 8.0E-05 |
| Positive regulation of myeloid leukocyte differentiation | 9 | 3.58 | 1.1E-04 |
| Response to low-density lipoprotein particle stimulus | 4 | 10.34 | 1.3E-04 |
| Regulation of cysteine-type endopeptidase activity | 25 | 2.03 | 2.3E-04 |
| Regulation of meiosis | 8 | 4.77 | 2.5E-04 |
| Positive regulation of behaviour | 16 | 2.73 | 3.2E-04 |
| Regulation of meiotic cell cycle | 8 | 4.00 | 5.2E-04 |
| Response to estrogen stimulus | 20 | 2.30 | 7.7E-04 |
| Positive regulation of histone acetylation | 6 | 5.82 | 1.1E-03 |
| Negative regulation of glycolysis | 4 | 12.41 | 1.4E-03 |

### Key myeloid differentiation transcription factors are differentially expressed

From analysis of the expression and methylation profiles of important myeloid differentiation regulatory transcriptions factors, it was observed that CEBPA (Supplementary Figure S3 A&B) and GFI1 (Supplementary Figure S3 C&D) may be required to maintain HL-60/S4 in the undifferentiated state (Figure 5). As such, downregulation of CEBPA is necessary for the further differentiation of HL-60/S4 to either the neutrophil-like or macrophage-like state. Meanwhile, SPI1 and CEBPB are upregulated in both differentiated states (Supplementary Figure S3 E&F and K&L).

**Figure 5:**
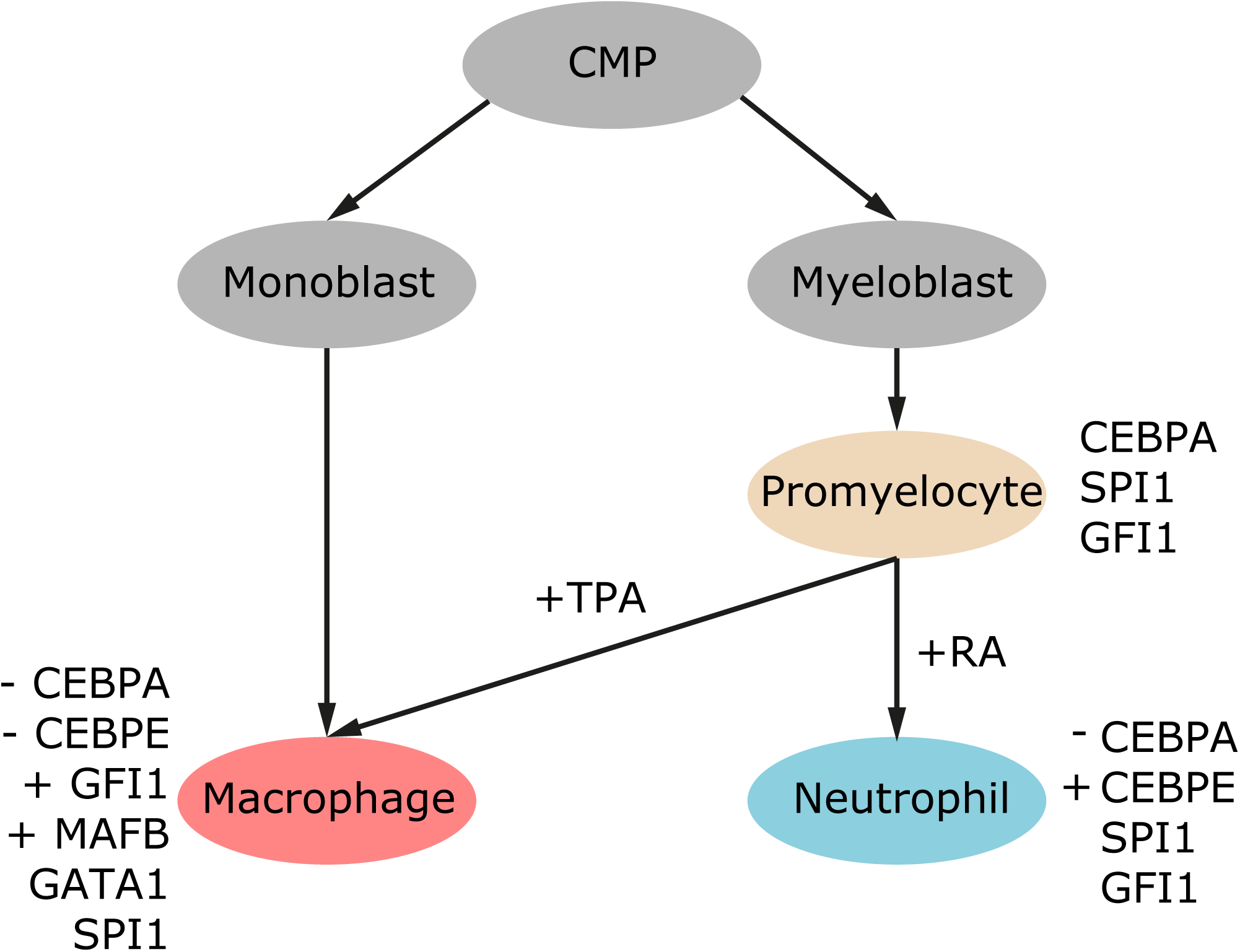
Chemical differentiation model of HL-60/S4 showing the transcription factors that may play an essential role in determining cell fate. Downregulation or upregulation of gene expression are denoted by “-“ or “+” respectively. Genes with no sign attach implies their levels are maintained at similar levels as in UN (promyelocytic) state.

Upregulation of CEBPE (Figure 4B) is seen in RA; whereas, it is downregulated in TPA, together with GFI1. In TPA treated cells, MAFB is upregulated, although still at low levels (Supplementary Figure S3 G&H). GATA1 is also down-regulated in RA and upregulated in TPA treated cells (Supplementary Figure S3 I&J).

## Discussion

### Differential methylation during HL-60/S4 differentiation occurs over small regions

Only small differences in DNA methylation were observed during HL-60/S4 cell differentiation at the 10 Mb window scale (Figure 1 E). Despite the lack of large scale methylation changes during the induced differentiation of HL-60/S4 cells, we observed both hyper-and hypo-methylation of a large number of differentially methylated single CpGs (DMPs) with a mean difference in methylation rate of 0.2 (Figure 2 A, C and D). Interestingly, the methylation rates of most of the differentially methylated CpGs ranged from around 0 to 0.4, corresponding to the partially methylated or unmethylated CpGs (Figure 2 C and D). This explains why only very few differentially methylated CpGs could be identified, since CpGs with this methylation rate value range were globally very sparse.

### The DNA methylation landscape of RA cells is closer to undifferentiated HL-60/S4 cells, than to TPA treated cells

Despite the generally similar megabase-scale methylation landscape observed in all 3 samples, they could be clearly distinguished using principal component analysis (Figure 1 C). Whereas TPA cells were seen to be very different from UN cells based on their whole genome methylation profiles, RA and UN cells were closely positioned on the axes of both principal component 1 and 2. Neutrophil methylation has already been shown to be only slightly, but significantly, different from the promyelocyte precursor cell methylation (Alvarez-Errico, et al., 2015). Thus, the small differences seen between RA (granulocyte-like) cells and the UN (promyelocytic) cell forms are consistent with that previous study.

### Differential methylation is limited to a few CpGs with very low levels of methylation

A total of 41,306 CpGs were identified to be differentially methylated; a very small number compared to the genome wide CpG numbers. The numbers of differentially methylated CpGs identified by a comparison of TPA to RA and UN cells were very similar. The lowest numbers of differences of differentially methylated CpGs were seen in a RA comparison to UN cells (Figure 2A). Hyper-methylated DMPs were only enriched in protein coding TSS and in CpGI and Enhancers for both RA and TPA, compared to UN. However, CTCF sites were only enriched in hyper-methylated DMPs in the TPA-UN comparison (Figure 2B). Hypo-methylated DMPs were seldom enriched for any particular genomic feature, except for CpGI, which was enriched in the RA-UN comparison, while enhancers showed enrichment within the TPA-UN comparison.

Changes in gene expression profiles, regulated by enhancers, may play a major role in the differentiation of macrophage-like cells. Enhancers stand out from other genomic features for TPA differentiated cells, which are quite different from UN cells (Figure 1C). The DMP module (M6), which has full methylation of CpGs in UN and RA, but hypo-methylation in TPA, is the same module that shows the highest enhancer enrichment (Figure 2D&E). These observations emphasize the significance of hypo-methylation of enhancers in macrophage-like differentiation, as observed in TPA-treated cells. On the other hand, modules M1 and M4 which showed either hyper-and hypo-methylation, for RA compared to UN, showed little enrichment of any genomic features, except CpGI. This may suggest a fine tuning of expression for already active genes, while hypo-methylation of DMPs in module M6 hints at the activation of expression of genes that might not be expressed in UN or RA cells.

On a broader view, hypo-methylation of enhancers, epichromatin and chromatin interaction domains in TPA cells suggests a remodeling of the transcriptional regulatory circuits in this state, compared to the RA and UN cell states.

### Interplay of DNA CpG methylation and nucleosome occupancy is genomic context dependant

In TPA cells we observed lower nucleosome occupancy and hypo-methylation around the DMPs of module M6 (Figure 2E and Supplementary Figure S2). This module was also enriched for enhancers (Figure 2F). Similar observations were made for modules M7 and M12, albeit with lower levels of methylation change, enhancer enrichment and differential nucleosome occupancy changes. In modules M8 and M11, we observe hyper-methylation in TPA cells but no increase in nucleosome occupancy. These modules had little or no enrichment of enhancers. Similarly, other modules with hypo-methylation for either UN (modules M5 and M10) or RA cells (M8 and M9) did not exhibit reduced nucleosome occupancy, nor were they enriched in enhancers. This suggests that differential nucleosome occupancy that is associated with differential DNA methylation in our differentiation system occurs in the genomic context of enhancers. This is consistent with previous findings of changes of nucleosome occupancy and DNA methylation in regulatory genomics contexts of CTCF binding and promoters (Kelly *et al.*, 2012) during cellular differentiation.

### RA and TPA cells share only a few DMPs

We identified 12 clusters of DMP patterns, which we grouped into modules. These modules revealed that most of the identified CpGs were differentially methylated only in TPA cells, compared to the other differentiated states (Figure 2 E). The first 6 modules describe CpGs that were differentially methylated in one cell state, by comparison to one other cell state; while the latter 6 modules are for CpGs that were differentially methylated in one cell state, compared to the other two cell states.

Since the differentiation of HL-60/S4 into the granulocyte-like or macrophage-like state is a branched process and not linear, the effects of most of the CpGs that are differentially methylated in one direction may not be important to the other differentiation direction; unless, of course, the effect on CpGs is required for the differentiation process. It is conceivable that the effects of differentially methylated CpGs in modules M7-12 may be related to cell differentiation in general, while those in modules M1-6 may be related to specific developments of the different cell states.

### Both positive and negative correlations are observed comparing DNA methylation of TSS regions and levels of gene expression

Earlier reports suggested that methylation in the promoter and the first exon inversely correlated with gene expression (Brenet *et al.*, 2011; Jones, 2012). As such, it would be expected that in the HL-60/S4 cell differentiation system, DNA methylation in the TSS region of genes should correlate negatively with gene expression. However, we observed equal numbers of genes that showed either positive or negative correlation between TSS methylation and gene expression was about equal (Figure 3). This observation suggests that there are additional epigenetic modifications required at gene promoters to regulating transcriptional activity (Ford *et al.*, 2017) or that gene expression is determined by the epignenetic state of multiple regulatory elements and not just the promoter (Ong and Corces, 2011).

### Long-range chromatin interactions play an important role in HL-60/S4 differentiation

RNA expression of CEBPE exhibits a strong inverse correlation with differential methylation in a downstream region of the CEBPE gene. These regions have been shown to be interacting, employing CHIA-PET in the K562 leukemia cell line (Figure 4A) (Dunham *et al.*, 2012).

Similarly, a region within the promoter of PGP was identified to contain DMPs which correlated negatively with the RNA expression of CCNF (upstream of PGP) (Figure 4E and F). As these two genes transcribe in opposite directions, they may share the same promoter. However, this region, despite being in PGP, showed negative correlation with only CCNF. Being a Cyclin, it is involved in regulating the progress through the cell cycle, but the exact function in this process of differentiation is not clear. We have also presented evidence that methylation of chromatin interaction partners also plays a crucial role for expression of genes in HL-60/S4 cells (Figure 4).

### CEBPA downregulation and differential regulation of CEBPE expression are required of HL-60/S4 differentiation

TSS methylation and RNA expression of key myeloid differentiation transcription factors SPI1, CEBPB, CEBPE, CREBBP, CEBPA, DNMTs and HDACs were examined. CEBPA was observed to be hyper-methylated in RA and TPA compared to UN cells (Figure S3). This resulted in significant downregulation of expression of CEBPA in the differentiated states compared to UN cells.

SPI1 and CEBPA, together with CEBPB are known to be required for the maintenance of CMP and GMP developmental stages of myeloid cells (Alvarez-Errico, et al., 2015). However, it is the counter-interaction between SPI1 and CEBPA transcription factors that decides whether a GMP differentiates or not (Iwasaki & Akashi, 2007), since CEBPA is known to repress macrophage differentiation induced by SPI1. However, down-regulation of CEBPA expression in both RA and TPA suggests that it is significant in maintaining HL-60/S4 in the promyelocytic state. Thus, down-regulating CEBPA is key to macrophage differentiation; whereas, SPI1 is also expressed over 1.5-fold in both RA and TPA compared to UN cells.

Most of the other transcription factors necessary for the differentiation of macrophage and granulocytes are equally regulated by RA and TPA. An exception is CEBPE, which is upregulated in RA, but downregulated in TPA (Figure 4 A and B). This suggests that it is the downregulation of CEBPE which permits the differentiation of HL-60/S4 into the macrophage-like state.

Employing these observations, together with the data of Supplementary Figure S3, we have developed a model of the HL-60/S4 differentiation program based upon the transcription factors that may be required (Figure 5). In this model, we propose that down-regulation of CEBPA is necessary for differentiation of HL-60/S4 cells. Whereas, CEBPE is upregulated in the neutrophil-like state, its downregulation and the simultaneous upregulation of MAFB and GATA1 are necessary of macrophage-like differentiation. This supports the idea that CEBPE is necessary for the commitment of HL-60/S4 cells to a neutrophil-like state.

The upregulation in the expression of GATA1 and MAFB genes supports their role in committing HL-60/S4 cells to a macrophage-like state. We, therefore, postulate that HL-60/S4 cells may only differentiate into a macrophage-like state upon down-regulation of CEBPA, in the absence of CEBPE.

## Conclusions

The HL-60/S4 cell line is an excellent model system for myeloid leukemia and for cell differentiation studies, due to the capability of differentiating the (undifferentiated) promyelocytic cell line into macrophage-like and granulocyte-like states, following TPA and RA treatments, respectively. The 3 different states of this cell line show very high methylation levels for most CpGs, leaving only a few partially methylated or unmethylated CpGs. Genome wide DNA methylation analysis indicates that the methylation level of the granulocyte-like state differs only slightly from the undifferentiated form; whereas, the macrophage-like state is very different from the other two cell states.

We found 41,306 CpGs (of the ∼26.7×10^6^ measured CpGs) showed significant differential methylation upon differentiation of the HL-60/S4 cells, concentrated within a group characterized by very low to partially methylated CpGs. This is substantially fewer than the 4.93 million dynamic CpGs involved in B-cell maturation, most of which were found in later stages of differentiation (Kulis *et al.*, 2015). Furthermore, since differentiation into the macrophage-like and granulocyte-like states is a branched set of events, only a few differentially methylated CpGs are shared between the diverged cell states. Hence, most of differentially methylated CpGs are specific to either macrophage-like or granulocyte-like differentiation.

Similarly, differential methylation was limited to the genomic features that overlapped with CpGs that are not fully methylated. This explains why regulatory genomic features such enhancers, CpG islands and protein-coding gene TSS were enriched, while epichromatin was highly depleted in the differentially methylated regions. This could also imply that once a CpG becomes methylated, it is more likely to remain methylated, which is consistent with observations in previous studies (Senner *et al.*, 2012).

A gene encoding a key transcription factor in the differentiation of myeloid cells (CEBPA) was hyper-methylated in both RA and TPA treated cells. Hyper-methylation of the promoter of this gene, however, negatively correlated with gene expression, implying repression of transcription of CEBPA in both the macrophage-like and granulocyte-like states. CEBPE, on the other hand, was hyper-methylated and expression was down-regulated only in the macrophage-like cell forms. This implies that down-regulation of CEBPE is required for macrophage development. Experiments involving CEBPE “knockout/know-down” are required to examine whether down-regulation of CEBPA in HL-60/S4 cells will promote differentiation into a granulocyte-like state.

## Materials and Methods

### Samples

We used the human AML (acute myeloid leukemia) cell line HL-60/S4, available from ATCC (CRL-3306). Differentiation of this cell line was induced with retinoic acid (RA) and 12-O-tetradecanoylphorbol-13-acetate (TPA) to attain the granulocyte-like and macrophage-like states, as previously described (Mark Welch et al 2017). In previous publications (Mark Welch et al 2017, Teif et al 2017), the undifferentiated (UN) HL-60/S4 cells were denoted “0”. In the current study the same undifferentiated cells are denoted “UN”.

### Sequencing and library preparation

Whole genome bisulphite sequencing (WGBS) libraries were prepared for untreated (UN), RA, and TPA treated HL-60/S4 cells. Libraries were prepared using the Illumina TruSeq DNA Sample Preparation Kit v2-set A (Illumina Inc., San Diego, CA, USA) according to manufacture guidelines. After the adapters were ligated to the library, they were treated with bisulphite followed by PCR amplification. Sequencing was performed on the Illumina HiSeq 2000 using paired end mode with 101 cycles using standard Illumina protocols and the 200 cycle TruSeq SBS Kit v3 (Illumina Inc., San Diego, CA, USA).

### Read alignment and methylation calling with BSMAP

WGBS sequencing data were analysed using BsMAP (Xi and Li, 2009) and BisSNP packages. In brief, sequencing reads were adaptor-trimmed using CUTADAPT package (Martin, 2011), while read alignments were performed against the human reference genome (hg19 GRCh37 version hs37d5-lambda, 1000 Genomes) using the BsMAP-2.89 package with non-default parameter –v 8 (Xi & Li, 2009). Putative PCR duplicates were filtered using Picard (version 1.61(1094) MarkDuplicates (http://picard.sourceforge.net). Only properly paired or singleton reads with minimum mapping quality score of >=30 and bases with a Phred-scaled quality score of >=10 were considered in methylation calling using the BisulfiteGenotyper command. BisulfiteTableRecalibration was called with –maxQ 40. Methylation calling was done with BisSNP package (Liu, et al., 2012) and single-base-pair methylation rates (b-values) were determined by quantifying evidence for methylated (unconverted) and unmethylated (converted) cytosines at all CpG positions. Non-conversion rates were estimated using data from mitochondrion DNA (chrM). Only CpGs with coverage greater than or equal to 10x in all sample replicates were considered in downstream analysis.

### Differentially methylated CpGs calling

Fisher exact test with α = 0.05 was applied to all 17,233,911 CpGs individually to extract differentially methylated positions (DMPs).

### Principal component analysis (PCA)

Principal component analysis was done on all 17,233,911 CpGs using the princomp command in R.

### Genomic features analysis

We extracted genic features (intron, exons, intergenic regions, genes transcription start site (TSS)) together with 4D genomic interaction data from gencode v17 (Harrow *et al.*, 2012), CpG Island, Laminal Associated Domains (LADS) and RepeatMasker definitions from UCSC (Rosenbloom *et al.*, 2013). Using the start and end coordinates of the genes from Genecode17, TSS was defined as the region extending 2kb upstream and 1kb downstream the start of the gene. RepeatMaskers considered in the enrichment analysis are: DNA repeat elements (DNA), Long interspersed nuclear elements (LINE), Low complexity repeats, Long terminal repeats (LTR), Rolling Circle repeats (RC), RNA repeats (RNA, rRNA, scRNA, snRNA, srpRNA and tRNA), Satellite repeats, Simple repeats (micro-satellites) and Short interspersed nuclear elements (SINE). Enhancer were extracted from ENCODE (Dunham *et al.*, 2012), FANTOM5 (Andersson *et al.*, 2017) and Vista (Visel *et al.*, 2007). Coordinates of HL-60/S4 Epichromatin are described (Olins *et al.*, 2014).

### Enrichment analysis

Genomic feature and chromosome enrichment in the DMPs were estimated using the formula:

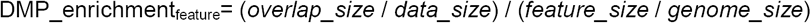

Where “*data_size*” is the size of the data (for either RA or TPA DMPs) been used to calculate the enrichment. Note that the enrichment of the hyper and hypo-methylated DMPs were calculated relative to the “*data_size*” or the total DMPs or DMRs called for each comparison but not relative to the total of only hyper or hypo-methylated DMPs or DMRs.

### Functional annotation

DMR functional annotations was performed with DAVID 6.8 (Huang, Sherman and Lempicki, 2009) using the full set of human genes as the background.

### Differential methylation patterns of DMPs analysis

DMPs were clustered using the hclust (Murtagh, 1985) with the complete linkage method after the Euclidean distances were calculated using the dist function in R. The hierarchically clustered DMPs were divided into 12 clusters using cutree. The resulting clusters were named as modules, from module M1 to module M12.

Feature enrichment within modules were estimated using the following formula:

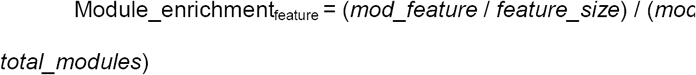

Where “*mod_feature*” is the size of a module overlapping with a specific genomic feature and *“feature_size”* is total size of a genomic feature in all modules. Whereas “*module_size*” is the total size of a module and “*total_modules*” is the size of the all modules together.

### Extraction of differentially methylated regions (DMRs)

DMR calling was done by first averaging coverage and number of methylation calls in a 3 CpGs sliding windows with maximum size of 2kb. Fisher exact testing was done using an alpha value of 0.05 to extract differentially methylated windows. Continuous differentially methylated windows were merged into one and Fisher test with same conditions were applied the second time ensure the regions were significantly differentially methylated. Differentially methylated regions that had 3 CpGs /1kb ratio were extracted before applying the final filter which states that a DMR should consist of at least 3 sliding windows. This step was to eliminate regions that probably had only one truly differentially methylated CpGs. As such, DMRs that were made of less than 3 windows (5 CpGs) were dropped also dropped.

### Differential gene expression

Differentially expressed genes data estimated using the RSEM software package (Li & Dewey, 2011) were obtained from our collaborators in The Josephine Bay Paul Center for Comparative Molecular Biology and Evolution (USA) (Mark Welch *et al.*, 2017).

### Correlation between gene expression and TSS methylation of HL-60/S4 genes

Methylation and transcriptome data were integrated by first extracting genes with –log2 (RNA expression fold change) >= 1.5 and TSS overlapping with at least one DMP as extracted using the Fisher exact test. Using this criterion we identified 114 and 221 genes for RA and TPA respectively, summing up to a total of 280 unique differentially expressed genes.

Secondly, genes with TSS overlapping with DMPs with methylation rate difference >=0.2 were extracted for functional annotation analysis. In this extraction criterion 86 and 112 genes were identified for RA and TPA respectively. The correlation between the average methylation change of DMPs overlapping with the TSS of a gene and the –log_2_ (RNA expression fold change) were estimated for RA and TPA genes in a scatter plot. Similarly, correlation between the average methylation of DMPs overlapping with a gene TSS and the gene’s expression were estimated using values from all samples and the distributions plotted separately for genes whose TSS overlap with RA and TPA DMPs.

Furthermore, the correlation between the methylation of individual CpGs in the gene body and TSS region and gene expression was estimated for all genes from both extraction criteria together with the gene expression of transcription factors known to be involved in myeloid cell differentiation (Figure 5).

## List of abbreviations

HL-60/S4: human myeloid leukemic cell line HL-60/S4 (ATCC CRL-3306).
UN: undifferentiated HL-60/S4
TPA: tetradecanoyl phorbol acetate treated HL-60/S4
CpG: Cytosine-phosphate-Guanine dinucleotide
CpGI: CpG island
DMP: differential methylated CpG position
DMR: differentially methylated CpG region
MPP: multipotent progenitor cells
CMP: common myeloid progenitor cells
GMP: granulocyte monocyte progenitor cells
MEP: megakaryocyte erythrocyte progenitor cell
WGBS: whole genome bisulphite sequencing
FISH: fluorescent in-situ hybridisation
M-FISH: multiplex FISH
LINE: long interspersed nuclear element
TSS: transcription start site
CHIA-PET: Chromatin interaction analysis by paired end tag sequencing
IM-PET: integrated method for predicted enhancer targets

## Author Contributions

DEO, RE conceived the research. RE, DEO, NI supervised the study. ALO, DEO acquired the samples and data. EBA, NI, VT processed the data. EBA, VT, MB, TB, ZG, NI analysed data. All authors interpreted and discussed data. EBA, NI wrote the paper. All authors commented on and critically revised the manuscript.

## Acknowledgements

We thank the DKFZ-Heidelberg Center for Personalized Oncology (DKFZ-HIPO) for technical support. We thank Anna Jauch for karyotyping the HL-60/S4 cells using M-FISH. We thank the College of Pharmacy (University of New England) for providing space and facilities to DEO and ALO, enabling the growth and characterization of HL-60/S4 cells.

## Competing interests

No competing interests declared

## Funding

ALO and DEO were Guest Scientists at the DKFZ Heidelberg, Germany) and recipients of support from the University of New England, College of Pharmacy.

## Data availability

Raw sequencing data was deposited at the ENA under accession PRJEB27665.

Associated processing scripts and differential methylation analysis scripts are available via GitHub: https://github.com/jokergoo/ngspipeline/blob/master/WGBS_pipeline.pl https://github.com/eantwibo/HL60S4_methylation_scripts/

## Supplementary figures

Supplementary figure S1: Genome of HL60/S4 is stable over long time and upon differentiation: Coverage plots of the WGBC data for UN (A), RA (B), and TPA (C) depicting stable genome during differentiation. D and E show 2 examples of M-FISH of undifferentiated HL-60/S4 over a period of 4 years depicting stability of the genome.

Supplementary figure S2: Average nucleosome occupancy around DMP of the different modules as described in figure 2. Each image shows nucleosome occupancy 2000 bases up-and downstream of DMPs per module. Nucleosome occupancy is shown in black, red and blue for untreated, RA and TPA treated respectively. GC content refers to the percentage of GC at each base relative to the DMP position.

Supplementary figure S3: Key myeloid differentiation transcription factors are differentially methylated and expressed during expression. DNA methylation landscape and gene expression of transcription factors know to play important role in myeloid differentiation. Gene expression levels for the three differentiation states as shown in the blue bar plots correspond with the methylation and correlation profiles on their left.

## Supplementary tables

**Supplementary table ST1:** Read and alignment statistics of the whole genome bisulphite sequencing data used in this study.

|  | UN | RA | TPA |
| --- | --- | --- | --- |
| QC-passed reads | 1,075,185,936 | 1,070,133,636 | 1,096,718,018 |
| Read pairs | 453,160,937 | 453,160,937 | 433,631,362 |
| Unpaired reads | 37,984,963 | 37,984,963 | 36,188,487 |
| Unmapped reads (%) | 12 | 10 | 18 |
| Duplicates (%) | 4 | 4 | 4 |
| Genome-wide coverage(x) | 28.87 | 29.43 | 27.56 |
| CpGs identified | 26681926 | 26699651 | 26647233 |
| CpG coverage | 21.9 | 22.6 | 20.2 |
| ChrM conversion | 0.998761 | 0.999071 | 0.998135 |

Supplementary tables ST1-ST13: enrichment of GO molecular function terms in modules M1-M12

[Submitted as an EXCEL file]

Supplementary table ST14: enrichment of GO biological process terms in module M6

[Submitted as an EXCEL file]

